# Omnitrophota dominate per-genome cross-replicon lateral gene transfer in a Fennoscandian deep-groundwater metagenome

**DOI:** 10.64898/2026.05.20.726742

**Authors:** Torben N. Nielsen

## Abstract

In a single Oxford Nanopore long-read metagenome from a Fennoscandian deep-groundwater borehole (KR0015B, Äspö Hard Rock Laboratory, Sweden), 791 protein clusters span at least one chromosomal contig and at least one co-sampled circular mobile element — the cross-replicon LGT-candidate cohort. The 199 participating chromosomes are dominated by three small-genome / symbiont-associated lineages — Patescibacteriota, Omnitrophota, and Nanobdellota — but the per-chromosome participation rate tells a different story: Omnitrophota chromosomes participate at an order-of-magnitude higher rate (mean 56 cross-replicon clusters per genome), while Patescibacteriota and Nanobdellota dominate by compositional abundance only.

Two large divergent circular mobile elements (233-kbp u20424375 and 123-kbp u29249220) — each lineage-restricted within a single Omnitrophota genus, with sparse cross-phylum reach (only 12 of their combined 289 cross-replicon clusters involve a non-Omnitrophota partner) — together account for 37% of the cohort and lack canonical plasmid or phage signatures. The 233-kbp element carries a Mu-class DDE transposase, is found integrated in one host chromosome at 99.3% nucleotide identity over 87% of element length, and carries an essentially complete bacterial big-operon r-protein cluster (31 r-protein KOs) as cargo with no rRNA genes — a cargo profile with no published precedent in the mobile-element literature. Seven cross-replicon clusters span both domains; per-cluster phylogenies confirm gene-tree topologies that violate the species-tree expectation in 6 of 10 callable smoking-gun trees. We release the cross-replicon cluster table, integrated mobile-partner classifications, and chromosome taxonomy as a community resource.

A parallel cross-chromosome catalog without the mobile-partner requirement contains 957 clusters, 95% of which carry no co-sampled circular plasmid or virus partner — a chromosome-only LGT footprint that bounds the MGE-coupled cohort and is consistent with vehicle-free / direct-contact transfer in lineages whose close-contact symbiotic biology is well-documented.

## Introduction

Lateral gene transfer (LGT) — the movement of genetic material between organisms outside vertical inheritance — is widely recognised as a major driver of microbial evolution (Soucy et al. 2015), but evidence for any individual transfer event is typically inferential. Phylogenetic-incongruence methods compare gene trees to species trees and flag mismatches as putative transfer events; these methods are powerful at scale but vulnerable to artifacts of incomplete lineage sorting, rate variation, and undersampling, and they say nothing about how the gene moved. Sequence-composition methods (codon usage, GC content, dinucleotide signatures) can localise recently-acquired regions to specific donors, but degrade rapidly as transferred sequence ameliorates to the recipient genome. The tightest lines of evidence link the same gene, at near-identical sequence, to both a host chromosome and a co-sampled mobile element (plasmid, virus, or integrative element) — placing the mobile vehicle and a chromosomal copy in the same physical sample at the same time. The criterion is high-specificity but not exhaustive: a recent transfer leaves a chromosome-shared signal without a co-sampled mobile partner whenever the vehicle has been lost, is non-circular, or never existed, such as in direct cell-to-cell contact.

Smillie et al. (2011) demonstrated this approach at scale on cultured genomes, identifying more than 10,000 recent transfer events along ecological rather than phylogenetic axes. Subsequent work has extended the framework to environmental metagenomes (Brito 2021, Camargo et al. 2024) but has been limited by the difficulty of recovering both chromosomal and mobile circular contigs from the same sample at scale. Long-read metagenomes have recently changed this constraint: Oxford Nanopore assembly of complex environmental samples now routinely yields hundreds to thousands of single-contig circular genomes per sample, including chromosome-scale and plasmid-scale elements in the same assembly.

The Äspö Hard Rock Laboratory provides a self-contained subsurface environment that has been the subject of metagenomic and viromic characterisation for over a decade (Wu et al. 2016, Holmfeldt et al. 2021, Mehrshad et al. 2021, Dopson et al. 2024). Holmfeldt et al. (2021) profiled the viral diversity of Äspö borehole samples but did not link viral genes to the bacterial and archaeal chromosomes assembled from the same boreholes; Dopson et al. (2024) characterised host genomes and biogeochemistry but did not couple the analysis to mobile elements. The cargo carried on these phages and on co-resident plasmids has not been catalogued at the scale or topology needed for sample-level LGT inference.

Here we apply a cross-replicon LGT-candidate filter to an Äspö borehole metagenome, KR0015B, in which Oxford Nanopore long-read sequencing recovered 9,382 circular contigs. We cluster the 1,082,188 predicted proteins from these contigs at high stringency (≥70% amino-acid identity over ≥80% mutual coverage) and retain those clusters that span both chromosomal and mobile contigs (the cross-replicon cohort), alongside a complementary catalog that drops the mobile-partner requirement to bound the chromosome-only LGT footprint (the cross-chromosome catalog). The biological questions are concrete: which lineages participate disproportionately in cross-replicon transfer, what kinds of mobile elements carry the cargo, how often does the cargo cross domain or phylum boundaries, and what fraction of the chromosome-only LGT signal lacks any co-sampled mobile vehicle?

## Results

### Recovery of 9,382 circular contigs and 791 cross-replicon LGT candidates

The KR0015B assembly produced 9,382 circular contigs ranging from 5,012 bp to 7.6 Mbp. 237 met the high-quality complete-genome threshold and define the chromosome set; the remainder are candidate mobile partners. After high-stringency protein clustering and exclusion of universal housekeeping markers, 791 clusters span at least one chromosome and at least one mobile partner, drawing on 199 chromosomes and 367 mobile contigs (Figure 1). Of those, 40 cross two or more phyla on the chromosomal side — 16 of them three or more — and 7 cross both domains.

**Figure 1.**
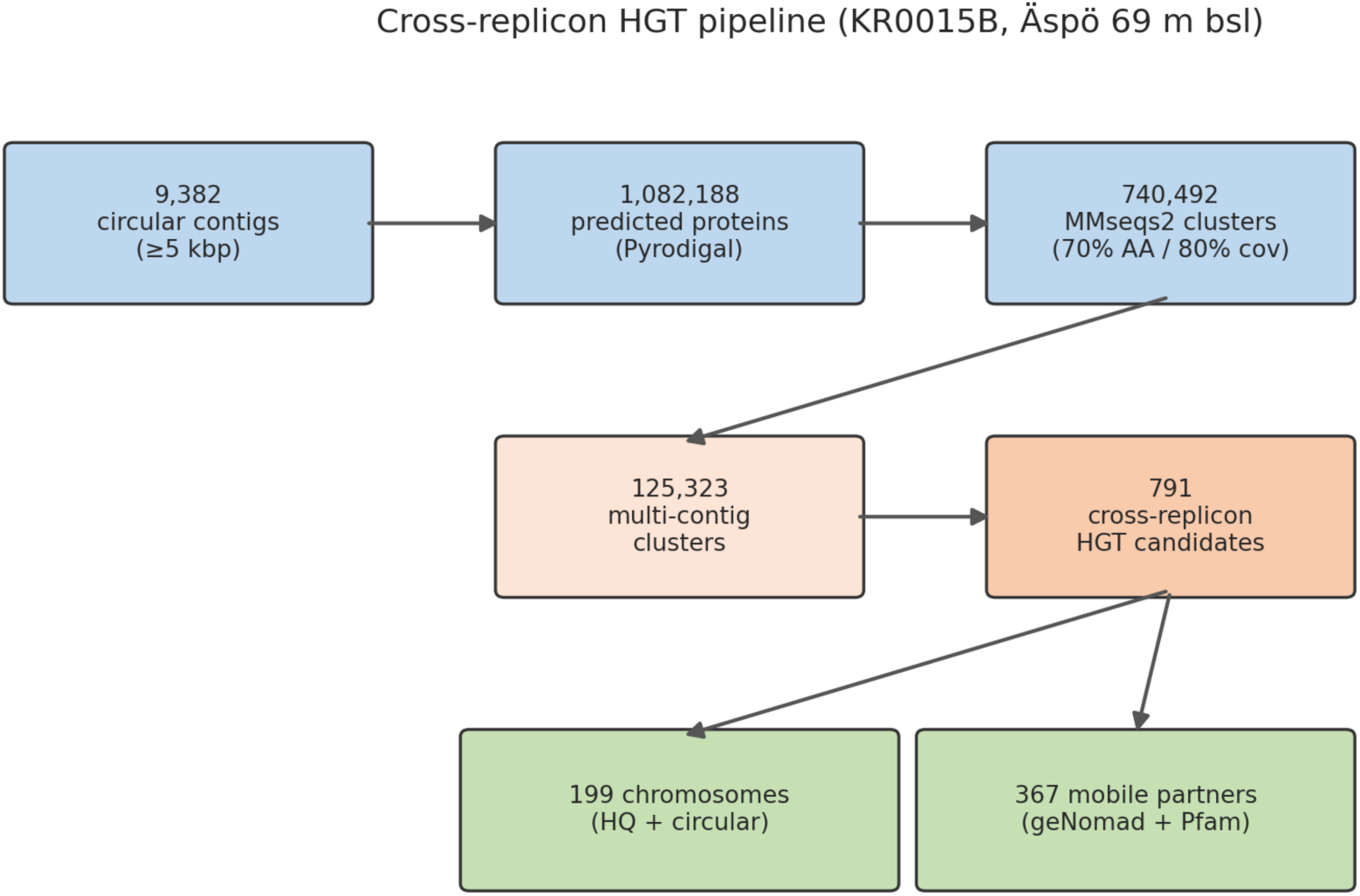
Pipeline schematic. Single-sample input (9,382 circular contigs ≥5 kbp from KR0015B Oxford Nanopore assembly) → Pyrodigal-predicted proteins → MMseqs2 70%/80% clusters → cross-replicon filter → housekeeping-marker exclusion → 791 cross-replicon LGT candidates spanning 199 chromosomes and 367 mobile partners.

### Patescibacteriota, Omnitrophota, and Nanobdellota dominate the participating chromosomes

The 199 participating chromosomes split 168 bacterial and 31 archaeal under GTDB-Tk classification. The bacterial side is heavily skewed toward two phyla:

| Bacterial phylum | Chromosomes |
| --- | --- |
| Omnitrophota | 77 |
| Patescibacteriota | 71 |
| Desulfobacterota | 4 |
| Planctomycetota | 4 |
| Campylobacterota | 3 |
| Auribacterota | 2 |
| Bacteroidota | 2 |
| CAIJMQ01 | 2 |
| CAYTYR01 | 1 |
| Chloroflexota | 1 |
| Spirochaetota | 1 |

Patescibacteriota and Omnitrophota together account for 148 of 168 bacterial chromosomes in the cross-replicon cohort. Both lineages are well-characterised as small-genome, often host-associated bacteria (Castelle et al. 2018, Probst et al. 2018).

The archaeal distribution is similarly concentrated:

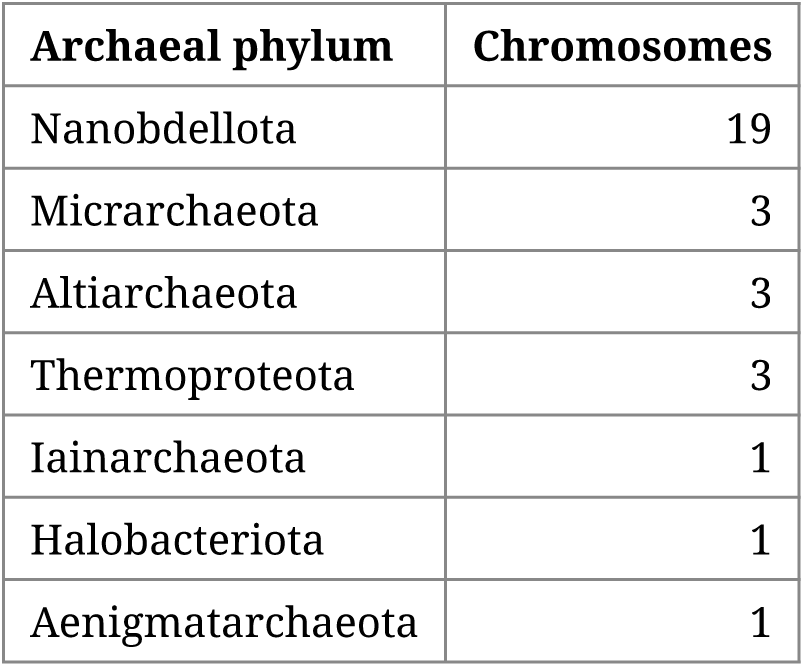

19 of 31 archaeal chromosomes are Nanobdellota — a DPANN phylum. Combined with the bacterial result, 167 of 199 participating chromosomes belong to small-genome / symbiont-associated lineages (Figure 2). The compositional concentration is qualitatively consistent with prior reports of mobile-element activity in DPANN and Patescibacteriota (Castelle et al. 2018, Wu et al. 2024) but, to our knowledge, has not been quantified at scale on a single environmental sample.

**Figure 2.**
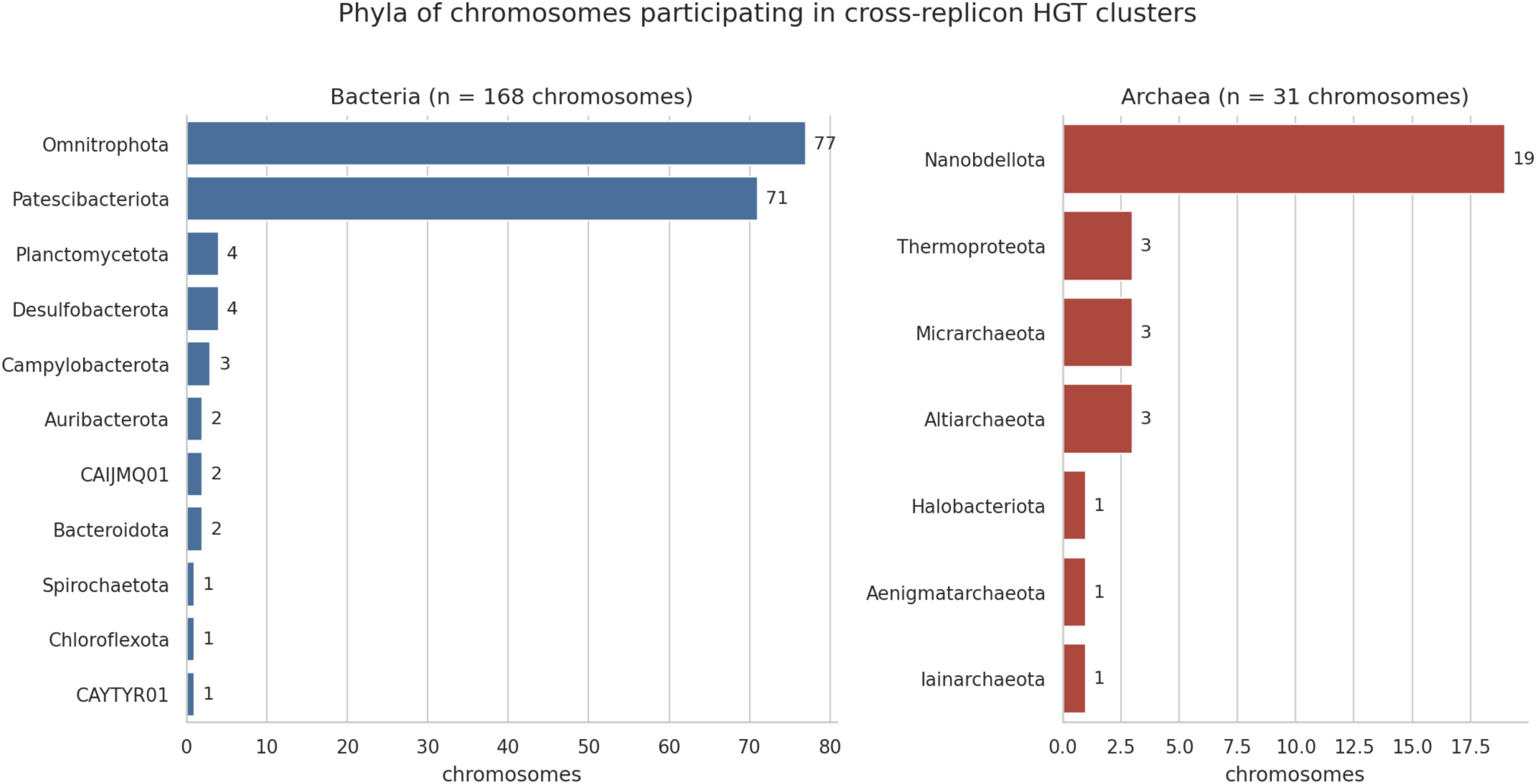
Phylum distribution of the 199 classified chromosomes participating in cross-replicon LGT clusters. Bacteria (left) are dominated by Patescibacteriota and Omnitrophota; Archaea (right) are dominated by Nanobdellota. The cross-replicon participant cohort is heavily skewed toward small-genome / symbiont lineages.

### Per-chromosome cross-replicon participation rates differ sharply by phylum

The compositional dominance of the three phyla decomposes unevenly. The 77 Omnitrophota chromosomes carry 4,347 of 4,937 cross-replicon cluster-memberships in the 199-chromosome universe — a mean of 56 clusters per genome, the largest per-genome signal in the assembly among phyla with more than three participating chromosomes. Patescibacteriota (mean 2.3 clusters per genome) and Nanobdellota (mean 1.8), by contrast, participate at or below the assembly mean: their dominance in the participant cohort reflects abundance, not an elevated per-chromosome rate. Per-phylum rate statistics are in Supplementary Table S7.

### A 233-kbp Mu-class circular element with one verified Omnitrophota integration host

The largest cross-replicon footprint of any single mobile contig belongs to u20424375, a 233,228-bp closed circular element at 43× read coverage, classified plasmid (single line of evidence) by the integrated scheme (one Pfam plasmid HMM, no phage hits, geNomad unresolved). It shares at least one cross-replicon protein cluster with 92 distinct chromosomes across 8 named bacterial phyla, but the bulk of its cluster sharing is heavily concentrated within a single Omnitrophota genus:

| Partner phylum | Chromosomes |
| --- | --- |
| Omnitrophota | 77 |
| Planctomycetota | 4 |
| Campylobacterota | 3 |
| Auribacterota | 2 |
| CAIJMQ01 | 2 |
| Desulfobacterota | 2 |
| CAYTYR01, Spirochaetota | 1 each |

The element participates in 195 of 791 cross-replicon clusters. Each cluster is a distinct protein family containing at least one u20424375 member, so at minimum 195 of the element’s ∼208 predicted proteins are part of the cross-replicon cohort — essentially the entire element behaves as cargo, not a small accessory subset.

The sharing pattern decomposes by partner taxonomy. Of the 195 clusters, 111 (57%) involve only same-genus partners within the JAHKAP01 genus of Omnitrophota (chromosomes u17976357, u4770637, u26960546); 77 (40%) reach other Omnitrophota genera; and only 7 (4%) involve a non-Omnitrophota partner. The “8 phyla” partner span is therefore a long-tailed signature: the bulk of cross-replicon clustering is within one Omnitrophota genus, with a smaller phylum-internal spread to other Omnitrophota and a small handful of cross-phylum reaches. This is the host-range signature expected for a large mobile element whose replication and integration machinery requires host-specific tuning (Maddamsetti et al. 2025, Smillie et al. 2010): copy-number control via Rep / iterons, partition systems coordinating with host nucleoid architecture, and integrase chemistry tuned to specific att sites all constrain large-element host ranges to closely related lineages. The cross-phylum reach is consistent with cargo-protein-mediated cluster formation (universal-conservation cargo proteins clustering across distant chromosomes) rather than active element-mediated transfer between phyla.

The cargo block delivered to each partner chromosome is not uniform; the distribution of shared-cluster counts across the 92 partners is bimodal:

| Shared protein clusters per partner | Number of partner chromosomes |
| --- | --- |
| 1 | 9 |
| 2–4 | 8 |
| 5–9 | 7 |
| 10–29 | 28 |
| 30–99 | 37 |
| ≥100 | 3 |

Median 20 clusters per partner; max 187. The three top-tier partners are the JAHKAP01 trio (u17976357, u26960546, u4770637), which share 187, 169, and 156 clusters respectively. Whole-genome alignment (minimap2 `-x asm5`) of u20424375 against all 237 KR0015B chromosomes recovers exactly one recent integration host: u17976357, in which 87% of the element aligns at 99.3% nucleotide identity over chromosomal positions 474,902–704,362. The other two JAHKAP01 partners do not carry an aligned integrated copy at asm5 stringency; their high cluster overlap reflects shared chromosomal content — same-genus orthology of the operons that u20424375 carries — rather than independent integrations of the element. Most partners (65 of 92) share 10–99 clusters, indicating substantial multi-protein cargo blocks; the 9 single-cluster partners are peripheral recipients of individual accessory genes. The host-range network is shown in Figure 4.

The cargo content of u20424375 is unusual. KofamScan annotation against the canonical KEGG r-protein KO panel recovers 31 r-protein KOs on the 233-kb element (19 large-subunit, 12 small-subunit), arranged in canonical bacterial big-operon order: the S10/spc/α superoperon at hub positions 64–73 kbp, the *rplKAJL* operon at 49–52 kbp, and the *rpsM-rpsK-rpsD* operon at 76–78 kbp, plus L17, L32, S16, EF-G, IF-1, alanyl-tRNA synthetase and three tRNA-modification enzymes elsewhere on the element. These positions correspond to the integrated copy’s chromosomal location at 635–664 kbp on u17976357. 26 of these r-proteins lack a detectable native paralog elsewhere on u17976357’s chromosome — they are u17976357’s only chromosomal copy of those proteins, mobilised in episomal form by the element. The episomal contig also carries no rRNA genes (cmsearch against bacterial and archaeal SSU/LSU/5S Rfam covariance models recovers zero hits) and only four tRNA genes (Thr, Trp, Ile2, Val). The cargo profile therefore differs from giant-virus translation cargoes (e.g., Tupanvirus and Klosneuvirinae carry many tRNAs and aminoacyl-tRNA synthetases but no ribosomal proteins; Abrahão et al. 2018, Schulz et al. 2017): r-protein-rich, rRNA-devoid, tRNA-poor.

Hub position 119,704–120,768 carries a Mu-class DDE transposase (Pfam Transposase_mut, E = 1.9 × 10⁻⁸⁵) that lies in the ∼2.6 kb arc of the circular element (hub positions 118,229–120,858) absent from the chromosomal integrated copy — the att region at the boundary between integrated and excised forms. No tyrosine recombinase (Phage_integrase) or serine resolvase / integrase Pfam profile hits the element at any threshold. The element’s mobility apparatus is therefore replicative-transposition-class (Mu-like), not the tyrosine-recombinase-mediated integrase chemistry of canonical conjugative ICEs. The integration’s chromosomal flanks lack the 5-bp target-site duplication characteristic of canonical Mu (Symonds et al. 1987), but the element’s right boundary carries three tandem 12-bp repeats (TTGACTCAATAC × 3) consistent with a transposase-binding terminus. We accordingly describe u20424375 as an integrative mobile element of unclear class with Mu-class DDE transposase machinery, rather than as a canonical ICE.

A second large divergent element, u29249220 (123,247 bp closed circular; 195× read coverage; 94 cross-replicon clusters across 68 partner chromosomes; integrated classification plasmid (single line of evidence)), shows the same lineage-restricted pattern with a different cargo distribution. Its top partner u2110955 (Omnitrophota / Gorgyraeia / UBA6249) is in 94 of 94 of the element’s clusters — a 100% cluster overlap signature consistent with u2110955 being the element’s primary host. The remaining partners are dominated by the same UBA6249 genus (4 additional chromosomes accounting for 65 of 94 clusters between them and u2110955), with 24 clusters reaching other Omnitrophota genera and only 5 reaching beyond Omnitrophota. The element carries a discrete 8.7-kbp integrative module at positions 103,765–112,499 — a tyrosine integrase (HTH_23 + integrase catalytic domain), paired KilA-N / ORF6N regulatory proteins, two AAA-class ATPases, and accessory cargo (CDP-paratose 2-epimerase for LPS/O-antigen biosynthesis, cardiolipin synthase) — an integrative cargo block embedded within the larger element.

Together the two hubs account for 289 of 791 cross-replicon clusters. We screened both for chimerism and found no evidence of misassembly in either (Methods; Supplementary Table S8).

### Mobile partners are predominantly virus and plasmid in roughly equal proportions

We classified the 367 mobile partners by combining geNomad (Camargo et al. 2024) probability scores with hits to two Pfam-A HMM panels — one for plasmid markers, one for phage markers — and assigned each contig to a strong, single-line-of-evidence, both, or unclassified category. The resulting distribution:

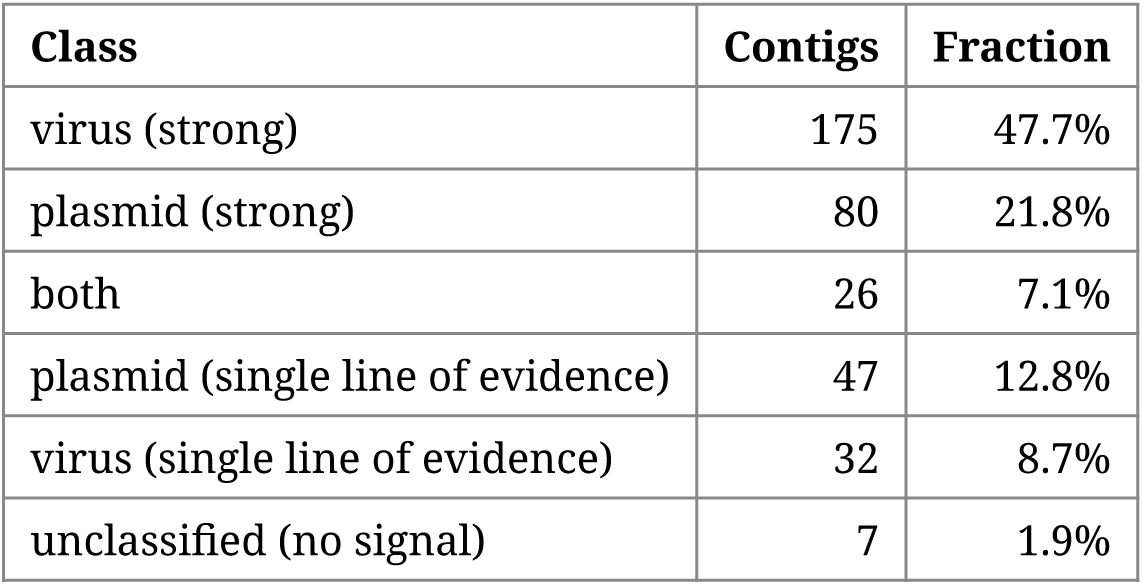

The plasmid fraction (22% strong + 13% single-line + a portion of `both`) is meaningfully higher than the equivalent fraction reported by geNomad alone, consistent with the Pfam panel surfacing plasmid-class evidence that geNomad’s neural-network model under-calls. On the contigs geNomad calls confidently — 52 plasmid, 182 virus — the integrated classification (or its weak / dual variants) preserves that call in essentially every case. The disagreement concentrates on the 133 contigs geNomad left unresolved: 77 of those were resolved as plasmids by the Pfam panel, 37 as viruses, 12 met the strong criteria for both panels, and 7 remain unclassified. This pattern is consistent with the lower plasmid recall reported by geNomad’s authors (plasmid MCC 0.778 vs virus MCC 0.953; Camargo et al. 2024). Figure 3 summarises the integrated classification across the 367-partner cohort.

**Figure 3.**
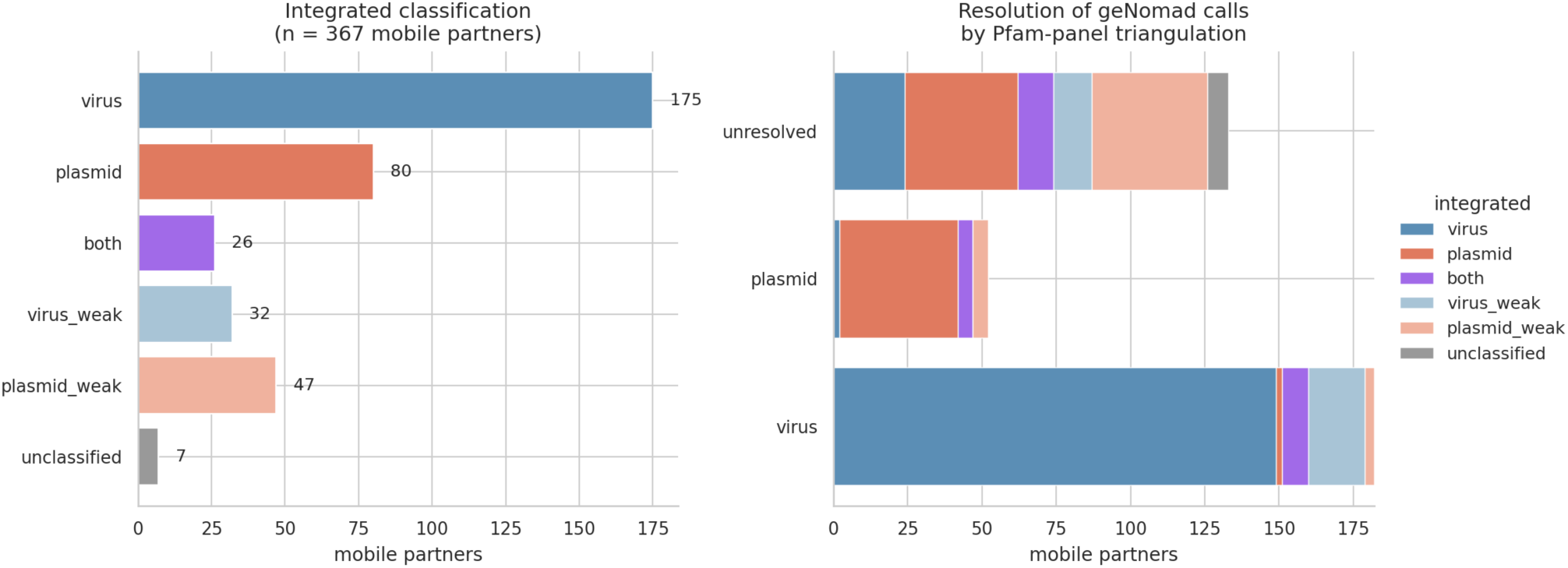
Mobile-partner integrated classification (n = 367). Left panel: counts per integrated class (virus, plasmid, both, virus single-line, plasmid single-line, unclassified). Right panel: how each geNomad call (virus / plasmid / unresolved) is partitioned by the integrated classification combining geNomad and Pfam-panel hits; the bulk of the geNomad-unresolved bar is recovered as plasmid by the Pfam panel.

**Figure 4.**
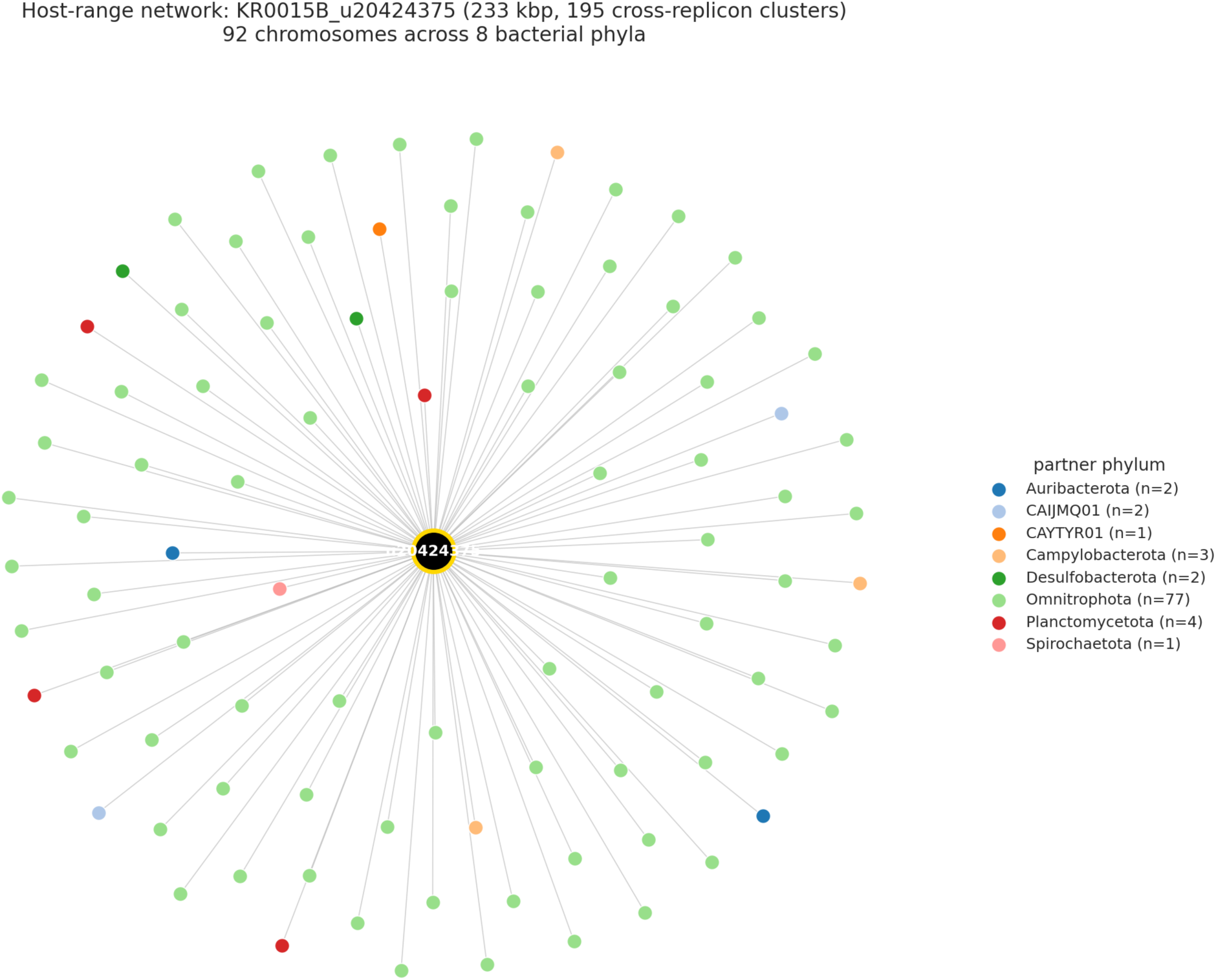
Large divergent integrative mobile element u20424375 (233 kbp closed circular, participating in 195 cross-replicon clusters). The 92 chromosomal partners are coloured by phylum; 8 distinct bacterial phyla are represented but partner-cluster counts concentrate sharply within Omnitrophota (and within the JAHKAP01 genus specifically). Edges connect the element to each chromosomal partner that shares at least one high-stringency protein cluster with it.

As a worked example of the plasmid (strong) end of the classification, u20671635 (99,932 bp circular) reads as a textbook conjugative plasmid by every signal: geNomad plasmid score 0.9988, 15 distinct Pfam plasmid HMMs matched (including MOB_Pre, Relaxase_3, TraI), and an essentially complete type IV secretion apparatus (T_virB2, T_virB5, T_virB8, T_virB9, T_virB10, T_virB11). Read coverage is a flat 50× across all three identity thresholds — a single dominant variant in the read pool, in contrast to the divergent hubs’ heterogeneous staircases. The integrated call is both because four Pfam profiles overlap the phage panel via integrase and integrative-element keywords; manual review confirms the dominant signal is plasmid. The contig appears in cross-replicon clusters with chromosomes spanning Patescibacteriota, Omnitrophota, and Planctomycetota, providing an independent positive control that the cross-replicon filter recovers genuine conjugative-plasmid–mediated cargo flow alongside the divergent-mobile-element signal.

### Confirmed Caudoviricetes phages account for a substantial fraction of cross-replicon signal

Across the full 367-partner cohort, virus partners (175 strong + 32 weak) outnumber plasmid partners (80 strong + 47 weak), reflecting a long tail of phages that participate in only a few clusters each. The picture inverts at the high end: among the top-50 mobile partners by cross-replicon involvement, the integrated calls are 18 plasmid-weak, 16 plasmid (strong), 6 both, 7 virus (strong), 2 virus-weak, and 1 unclassified — a plasmid-leaning distribution. The top two slots are occupied by the two large divergent integrative mobile elements characterised above; the remainder of the top-50 are dominated by plasmid-class contigs delivering cargo to multiple partner chromosomes.

The strong-virus partners all fall in Caudoviricetes (tailed double-stranded-DNA phages); the largest — jumbo phages of 156–258 kbp — carry the strongest hallmark enrichment scores (60–122×). This matches the prior virosphere characterisation of Äspö borehole samples (Holmfeldt et al. 2021), in which large dsDNA phages dominate the recoverable viral diversity.

### Cross-domain transfer events: archaea ↔ bacteria

Seven cross-replicon clusters span at least one archaeal and one bacterial chromosome. The largest:

- u16958799_186: 31 proteins, 23 chromosomes (8 archaeal + 15 bacterial) — the broadest cross-domain footprint in the cohort — and 1 mobile partner of 119 kbp.
- u27674409_1044: 19 proteins, 15 chromosomes (Thermoproteota + Omnitrophota), 1 mobile partner of 156 kbp.
- u7679499_281: 15 proteins, 11 chromosomes spanning 8 phyla (Auribacterota, Bacteroidota, Desulfobacterota, Halobacteriota, Omnitrophota, Patescibacteriota, Planctomycetota, Spirochaetota; Halobacteriota is the archaeal phylum), 3 mobile partners of up to 45 kbp.
- u33659312_8: 9 proteins, 6 chromosomes (5 archaeal: Micrarchaeota ×3, Iainarchaeota, Nanobdellota; 1 bacterial: Omnitrophota), 2 mobile partners of ∼12 kbp each. Both mobile partners are confidently called plasmids (geNomad plasmid score 0.9999 / 0.9998; 3 Pfam plasmid hits each). The cluster topology is shown in Figure 5.
- u14134137_29: 10 proteins, 7 chromosomes (1 Nanobdellota + 4 Patescibacteriota + 2 Omnitrophota), 2 mobile partners (∼31 kbp).
- u13132821_3: 5 proteins, 3 chromosomes (Iainarchaeota + Micrarchaeota + Omnitrophota), 2 ∼12-kbp plasmids.
- u31837952_2: 4 proteins, 2 chromosomes (Nanobdellota + Omnitrophota), 2 mobile partners (∼10–15 kbp) — the smallest cross-domain cluster and the simplest topology to interpret (one archaeal + one bacterial chromosome on plasmid-class mobile partners).

**Figure 5.**
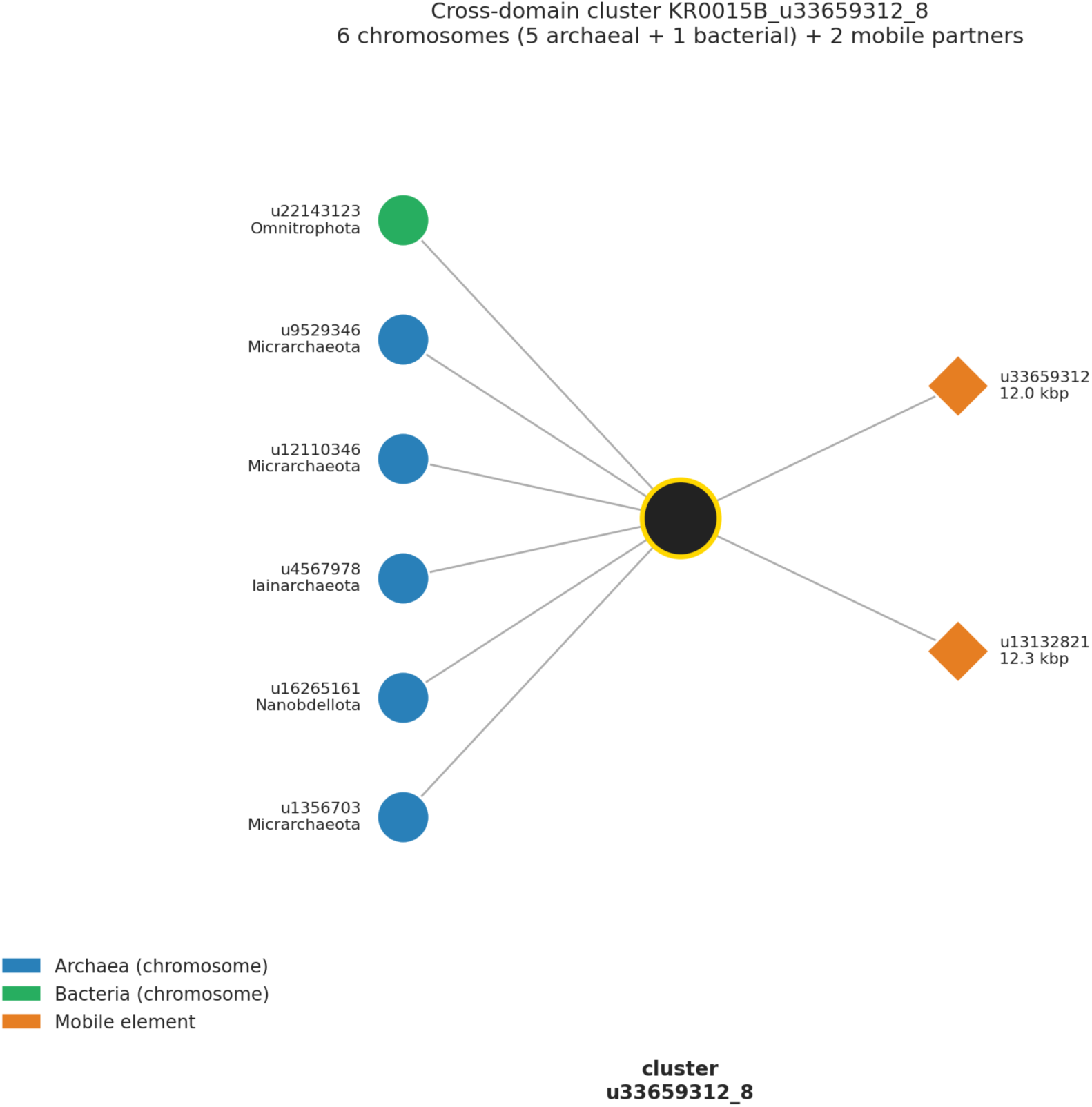
Cross-domain LGT cluster topology for u33659312_8 (9 proteins). Six chromosomal partners — five archaeal (Micrarchaeota ×3, Iainarchaeota, Nanobdellota; blue) plus one Omnitrophota (green) — share the cluster with two ∼12-kbp mobile elements (orange) classified as plasmids by both geNomad and the Pfam panel.

Across the seven, the archaeal participants concentrate in Nanobdellota (the DPANN superphylum) together with Micrarchaeota, Iainarchaeota, and Thermoproteota, and the bacterial participants concentrate in Patescibacteriota and Omnitrophota — the same lineages that dominate within-domain cross-replicon participation. Cross-domain LGT in this sample is therefore not a distinct phenomenon but a continuation of the same small-genome–to–mobile-element pattern stretched across a larger phylogenetic distance.

### Phylogenetic confirmation of LGT in a smoking-gun cohort

To distinguish genuine recent transfer events from orthology relics that happen to retain ≥70% AA identity across deep evolutionary distances, we built per-cluster maximum-likelihood phylogenies for a smoking-gun cohort of 12 clusters: the 7 cross-domain clusters above (≥1 archaeal + ≥1 bacterial chromosomal member) and the 5 cross-phylum clusters with the broadest phylum span (≥3 distinct bacterial phyla on participating chromosomes) that survive the housekeeping-marker filter. Two of the cross-domain clusters have only 2 or 3 leaves with assigned phylum and cannot return a non-zero phylum-disruption score by construction; they are listed in Table 1 for completeness but excluded from the callable evaluation. Member proteins were aligned with mafft L-INS-i, trimmed with BMGE (BLOSUM30) to retain phylogenetically-informative columns, and supplied to IQ-TREE 3 (Wong et al. 2026) with ultrafast bootstrap branch support.

**Table 1.**
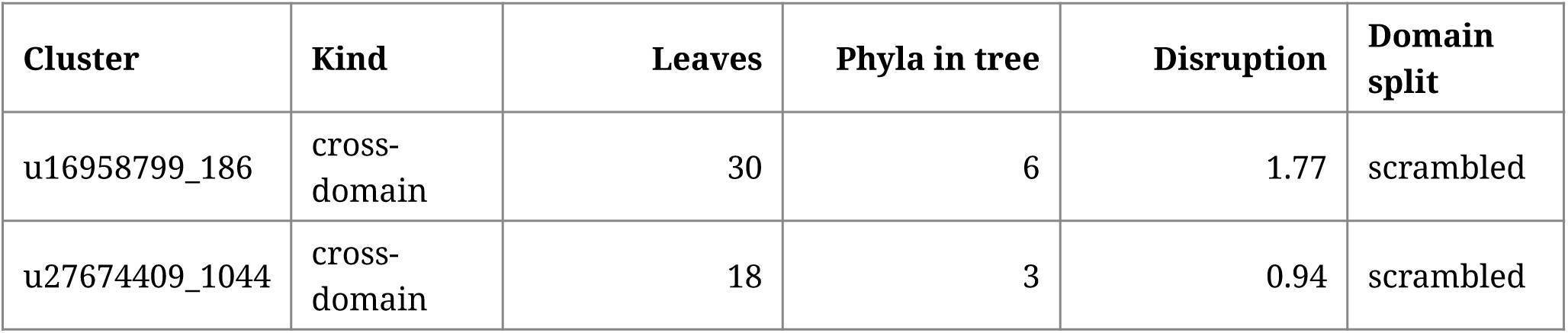
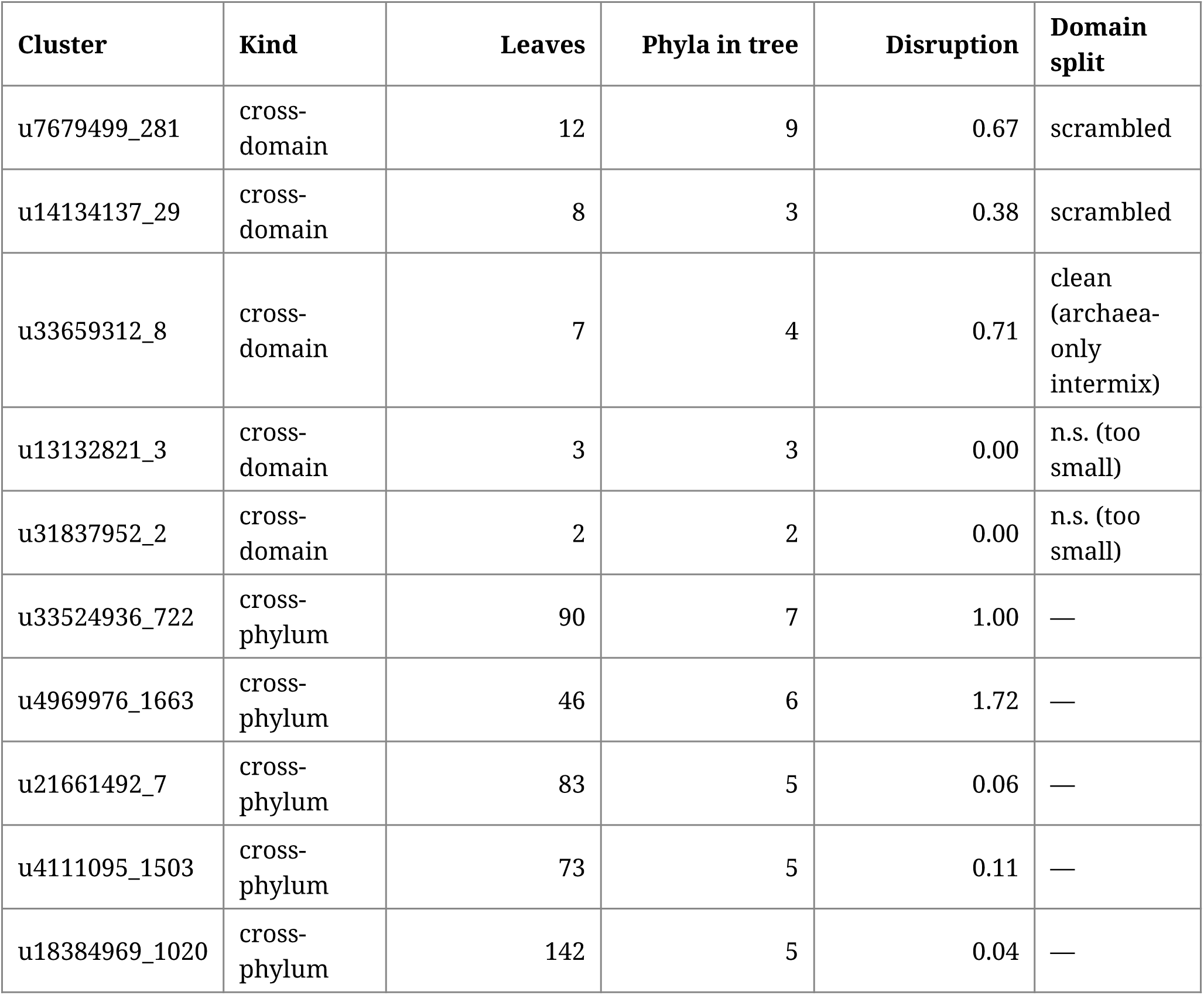
Phylum-disruption scores across the 12-cluster smoking-gun cohort.

For each tree we computed a *phylum-disruption score*: the sum, over each phylum present, of non-self leaves contained in the smallest clade encompassing that phylum’s members, divided by the total number of phylum-assigned leaves. A score of 0 indicates perfect phylum-monophyly (no LGT signal in the gene tree); scores ≥1 indicate that members of multiple foreign phyla intersperse with each phylum’s leaves to the point where every leaf has on average at least one foreign-phylum near-neighbour — a strong LGT signature. We additionally flagged whether archaeal and bacterial leaves form clean (sister-clade) splits or are intermixed in the cross-domain trees (Table 1).

Of the 10 trees with enough leaves to call (≥4 leaves with assigned phylum), 6 yield disruption scores ≥0.4, and 4 of the 5 callable cross-domain trees show non-clean domain separation — archaeal and bacterial leaves interspersed across the gene tree rather than splitting into reciprocal clades. The highest-disruption cross-phylum cluster, u4969976_1663 (46 leaves, 6 phyla), reaches a score of 1.72 with members of multiple bacterial phyla intermixed throughout the tree — the strong-LGT signature. Three other cross-phylum trees (u21661492_7, u4111095_1503, u18384969_1020) show low disruption (≤0.11) and group their members predominantly along phylum lines: cross-phylum *membership* alone is therefore insufficient to claim LGT, and phylogenetic confirmation is needed to discriminate. Across the cohort, gene-tree topology violates the species-tree expectation in the majority of callable cases, and the cross-replicon cluster signal acts as a faithful proxy for recent LGT in those clusters (Figure 6).

**Figure 6.**
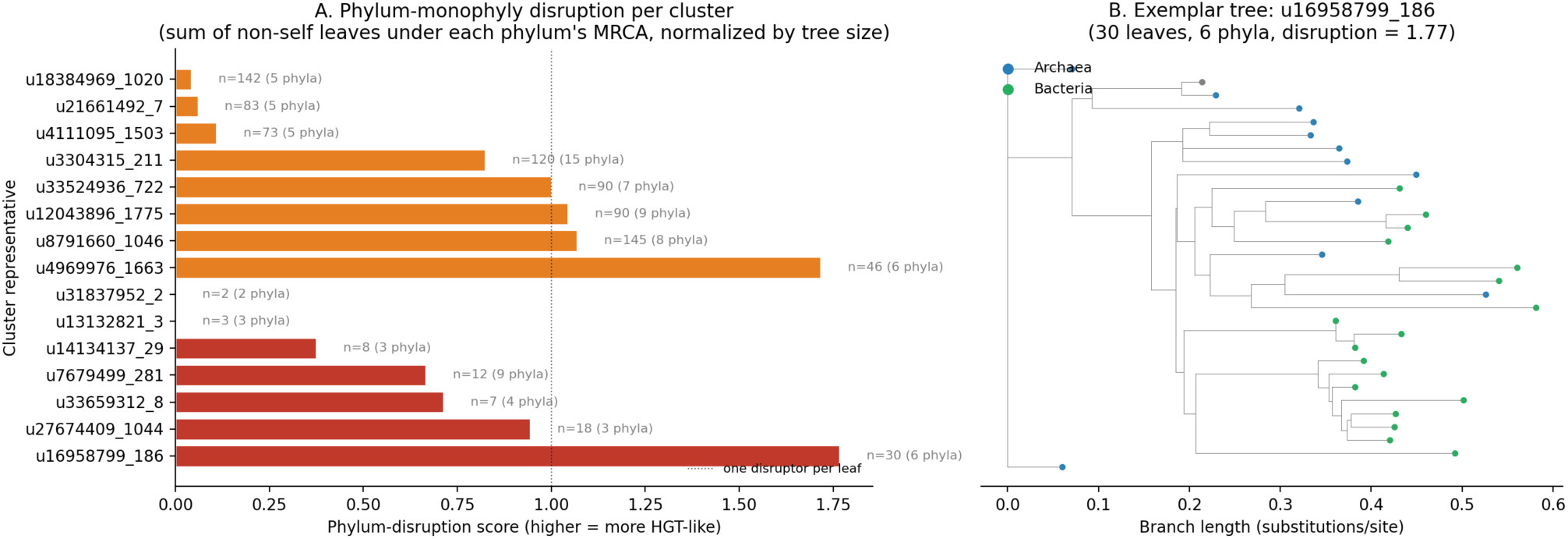
Smoking-gun cluster phylogeny disruption. Left panel: phylum-disruption score per cluster (red = cross-domain, orange = cross-phylum) with leaf and phylum counts; high disruption tracks the degree of phylum-leaf intermixing in the gene tree rather than phylum count alone — three cross-phylum clusters with ≥5 phyla nonetheless score ≤0.11 because their leaves group cleanly along phylum lines. Right panel: an exemplar protein tree with leaves coloured by domain (blue = Archaea, green = Bacteria), illustrating the interspersed topology that produces the disruption signal.

The full per-tree topology table (including per-phylum-disruption breakdowns and the leaf metadata) is released as Supplementary Table S4; cluster trees are at data/trees/<REP>/<REP>.treefile .

### Cross-chromosome LGT extends well beyond the cross-replicon cohort

The 791 cross-replicon cohort is conservative by design: each cluster must contain at least one chromosome and at least one circular mobile element clustered at high stringency, and clusters whose representative is a universal housekeeping marker (GTDB-Tk bac120 / ar53 single-copy panel) are excluded — the deep across-phylum conservation of these proteins makes their cross-replicon clustering uninformative about recent transfer. To estimate how much LGT this filter excludes, we built a parallel cross-chromosome catalog with the mobile-partner requirement dropped, retaining clusters whose members reach ≥2 complete-genome chromosomes from ≥2 distinct phyla and applying the same housekeeping filter.

The wider catalog comprises 957 clusters — 161 cross-domain (archaea ↔ bacteria) and 796 cross-phylum (within domain, ≥2 phyla). Only 47 of them overlap with the 791 cross-replicon cohort; the other 910 cross-chromosome LGT clusters carry no detectable circular mobile partner in the same sample. The largest cross-domain cluster in the wider catalog (u18384969_1839, 78 members, 71 chromosomes from 8 phyla on both sides of the archaea-bacteria divide) is invisible to the cross-replicon filter for this reason. The Venn-diagram overlap of the two catalogs and their composition by LGT kind are summarised in Figure 7.

**Figure 7.**
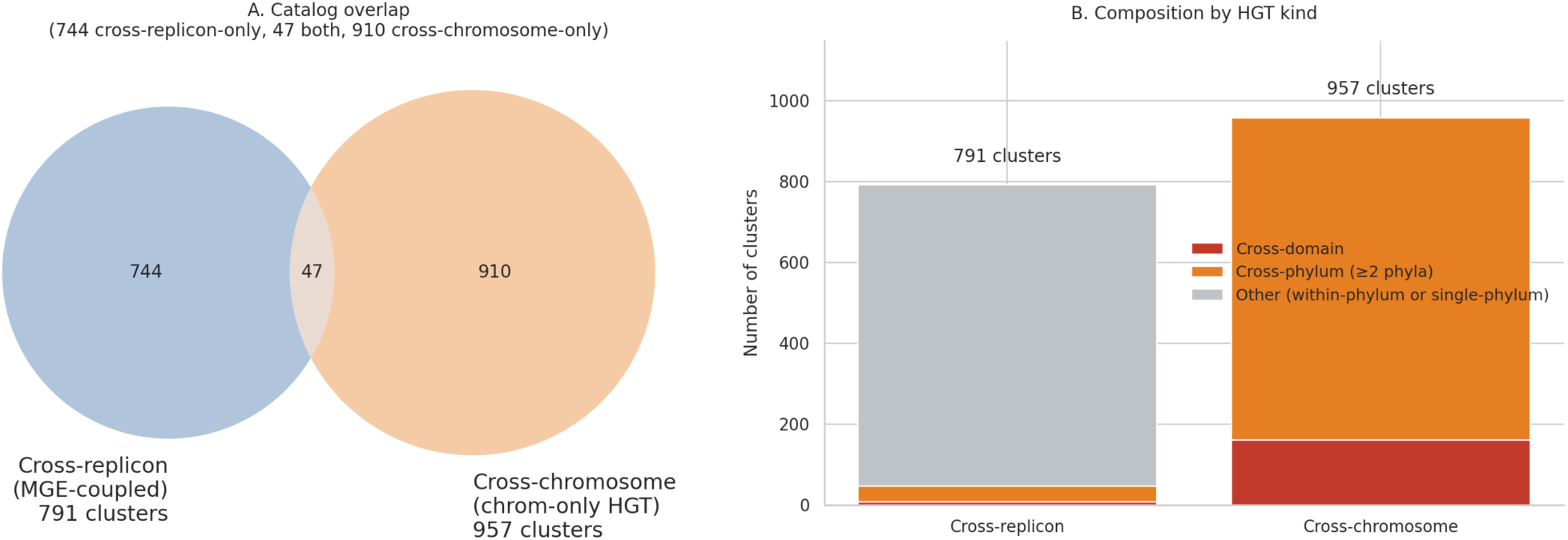
Dual-catalog comparison of cross-replicon (791 clusters; MGE-coupled LGT) versus cross-chromosome (957 clusters; chromosome-only LGT). **A**: Venn diagram of cluster overlap (47 shared; 744 unique to cross-replicon; 910 unique to cross-chromosome). **B**: composition by LGT kind — cross-domain, cross-phylum, other — for each catalog. The *rbcL* clusters from Nielsen and Lui (2026b) sit entirely on the chromosome-only side.

We read the 910 chromosome-only LGT signals as transfer events whose mobile vehicle either was not retained as a circular element in the sampled population, operates as a non-circular MGE outside our pipeline’s scope (transducing phages, gene transfer agents, integrated proviruses), or never passed through a discrete circular intermediate at all (the vehicle-free / direct-contact route is developed in the Discussion). The 791 cross-replicon cohort is therefore best understood as the MGE-coupled LGT cohort — a high-specificity subset in which the gene’s mobile vehicle is co-observed with its destination — and the 957-cluster catalog is the broader LGT footprint of which the cross-replicon set is a tractable, well-characterised slice. The full table is released as Supplementary Table S3.

### Functional annotation: cargo classes carried by cross-replicon clusters

We annotated the 791 cluster representatives against three independent sources: the Pfam-A HMM library, the KofamScan KEGG-ortholog (KO) panel (Aramaki et al. 2020), and UniRef90 via MMseqs2 translated search at minimum 30% identity and 50% query coverage. Coverage by source: 657 reps hit ≥1 Pfam-A family, 690 received a KO assignment, 709 hit a UniRef90 sequence, and 769 carry at least one annotation across the three sources (Figure 8A). The remaining 22 reps are completely dark by all three searches; 5 of them are members of clusters that include a hub-encoded protein (2 on u20424375, 3 on u29249220), the rest distributed across other contigs. The dark-rep cohort is small enough that its concentration on the hubs is at most a soft enrichment; the predominantly uncharacterised protein content of the two hubs is a separately observable property (see *u20424375* / *u29249220* descriptions in Results).

**Figure 8.**
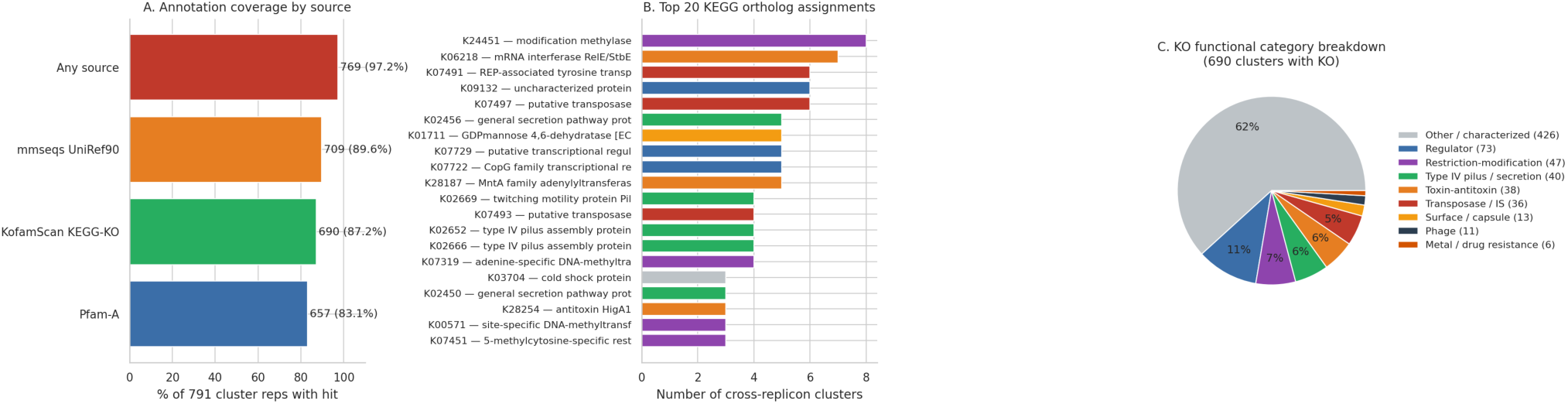
Functional annotation of the 791 cross-replicon cluster representatives. **A**: source-level coverage (Pfam-A, KofamScan KEGG-KO, mmseqs UniRef90, any source). **B**: top-20 KO assignments coloured by recognisable LGT-cargo category (transposase / IS, restriction-modification, toxin-antitoxin, type IV pilus / secretion, regulator, metal & drug resistance, surface / capsule, phage). **C**: KO-category composition across all KO-assigned clusters.

The 690 KO-assigned clusters partition by predicted function into the recognisable accessory-genome cargo classes that mobile elements carry between hosts (top-20 KOs in Figure 8B). Restriction-modification (47 clusters; modification methylases including K24451 and K07319, plus 5-methylcytosine-specific restriction enzyme A) and toxin-antitoxin systems (38 clusters; mRNA interferase RelE/StbE K06218, MntA K28187, HigA1/HigB1 K28253/K28254 pairs) — the canonical plasmid-maintenance armament — together account for 12% of the KO-assigned cohort. Insertion-sequence transposases account for 36 clusters (K07491, K07497, K07493, K07498, K07483, K18320), placing IS-element movement directly in the cross-replicon record. Type IV pilus and general secretion machinery (K02652 PilB, K02666 PilQ, K02669 PilT, K02456 secretion-G; 40 clusters across these and related KOs) places conjugation and twitching motility apparatus on cross-replicon mobile elements. Surface-modification gene families (LPS / capsule sugars; e.g., K01711 GDP-mannose 4,6-dehydratase, 13 clusters) and metal/drug resistance (CopG nickel-responsive regulator K07722, MerR copper efflux K11923, MFS DHA1 multidrug efflux K08153; 6 clusters) round out the recognisable mobile-cargo signal. The remaining 510 of the 690 KO-assigned clusters partition as 426 broadly distributed characterised protein families that do not match the cargo-category keyword rules, 73 general transcriptional regulators, and 11 phage structural / integration KOs (Figure 8C). The cross-replicon cohort is therefore dominated by cargo classes well-established as enriched on plasmids and integrative elements rather than by housekeeping genes — an independent functional cross-check that the filter selects for mobile-cargo proteins. The full per-cluster annotation table is released as Supplementary Table S6.

## Discussion

### Cross-replicon clustering as an MGE-coupled LGT filter

The cross-replicon filter is a coarse but interpretable LGT-candidate filter. A protein cluster that contains members on a chromosome and on a co-sampled mobile element places the gene’s mobile vehicle and a chromosomal copy in the same physical sample at the same time, and high-stringency clustering ensures that the chromosomal and mobile copies are too similar to represent ancestral orthology. Co-occurrence of a gene on a chromosomal contig and on a co-sampled mobile contig is symmetric: it does not by itself establish the direction of transfer, the donor, or the time of transfer. The filter is also conservative in that it requires the mobile vehicle to still be detectable in the sample — older transfers whose mobile precursor has been lost or has diverged below the clustering threshold will be missed. To our knowledge, comparable single-sample environmental catalogs of chromosome–MGE shared protein clusters have not been reported; published cross-replicon analyses have used pooled multi-sample (Pérez-Carrascal et al. 2025; Yu et al. 2024), short-read plasmidome (Kothari et al. 2019), or global cross-genome plasmid-network (Redondo-Salvo et al. 2020) designs, leaving the per-sample scale of cross-replicon protein sharing in environmental communities largely uncharacterised.

A second interpretive split is needed within the 791. Of those clusters, 47 reach ≥2 distinct phyla (40 cross-phylum within a single domain; 7 cross-domain) on the chromosomal side and are direct evidence of inter-host horizontal transfer — the same gene moving between chromosomes from different phyla, with its mobile vehicle co-observed in the sample. The remaining 744 clusters span chromosomes from a single phylum (or have only one chromosomal partner) plus a co-sampled mobile element, and are observationally compatible with two distinct mechanisms that the cross-replicon filter cannot separate: genuine inter-host transfer between two same-phylum hosts, or a chromosomal gene captured into the mobile pool of a single host (homologous recombination onto a co-resident plasmid, IS-mediated transposition, replication-time capture) without any inter-host event. The 47-cluster ≥2-phyla cohort is the conservative subset on which the strongest inter-host-transfer claims rest; the 744-cluster single-phylum cohort is the broader cross-replicon co-occurrence catalog, the contents of which include both confirmed and ambiguous transfer histories.

### Relationship to prior Äspö virosphere and biogeochemistry work

The Äspö Hard Rock Laboratory has been the subject of metagenomic and viromic characterisation for over a decade (Wu et al. 2016, Pedersen 2012, Holmfeldt et al. 2021, Westmeijer et al. 2022, Mehrshad et al. 2021, Dopson et al. 2024). Holmfeldt et al. (2021) characterised the Äspö virosphere as following slow-motion “boom and burst” cycles in which large viral populations periodically replace one another; the 175 strong-virus partners (and a further 32 single-line virus calls) we identify as cross-replicon partners in KR0015B are consistent with that framing in genus and class composition (Caudoviricetes-dominant, jumbo-phage-rich), and our cross-replicon cluster set provides the first systematic cargo-level link between those phages and the bacterial and archaeal chromosomes assembled from the same borehole. The biogeochemical and host-genome characterisation in Dopson et al. (2024) supplies the site context for KR0015B and is referenced here for ecological framing rather than directly compared.

### The chromosome-only-without-vehicle residual: three compatible mechanisms

The dual-catalog comparison reveals that 95% of cross-chromosome LGT clusters in this sample have no co-sampled mobile vehicle: 910 of the 957 cross-chromosome clusters do not cluster with any circular plasmid or virus contig at high stringency. Three routes are consistent with this pattern: the mobile vehicle may have been lost or diverged below the clustering threshold; the transfer may have used an MGE class our pipeline does not capture (transducing phages, gene transfer agents, integrated proviruses with no detectable circular form); or the transfer may never have used a discrete circular vehicle at all. Our data cannot apportion the 910 clusters among these three routes.

The third route, vehicle-free direct contact, is increasingly well-documented in the very lineages that dominate our cohort. Intercellular nanotubes (Dubey & Ben-Yehuda 2011) and DNA-bearing membrane vesicles (Renelli et al. 2004) move chromosomal DNA between cells without an intermediate circular element. Cryo-electron tomography of DPANN co-cultures (Johnson et al. 2024) and direct host-association data on Antarctic Nanohaloarchaea (Hamm et al. 2019) place this route firmly within the lifestyles documented for Patescibacteriota and DPANN. The high prevalence of host-attachment gene clusters in Omnitrophota (Seymour et al. 2023) — the per-genome cross-replicon hot-spot we identify above — together with the close-contact predator/symbiont biology of Patescibacteriota and Nanobdellota, give cellular geometry consistent with vehicle-free transfer in this sample. The present data do not measure how much of the 910-cluster residual any one of the three routes accounts for.

The 791 cross-replicon cohort and the 910 chromosome-only-without-vehicle cohort therefore sample different observable categories of LGT in the same metagenome. The 791 are recent inter-host transfers that left a co-sampled circular vehicle. The 910 are the broader chromosome-only footprint, whose mechanistic apportionment among the three routes above is a question this sample alone cannot resolve.

### Small-genome lineages dominate the cross-replicon participant set

Patescibacteriota, Omnitrophota, and Nanobdellota together account for 84% of the 199 classified chromosomes participating in cross-replicon clusters, but the per-chromosome participation rates pull these three lineages apart. Omnitrophota chromosomes participate in cross-replicon clusters at roughly an order of magnitude higher rate than the rest of the chromosome universe (mean 56 clusters per genome), and the 77 Omnitrophota chromosomes alone carry 88% of all cross-replicon cluster-memberships in the sample. This is the genuine per-genome enrichment in the cohort. Patescibacteriota and Nanobdellota, by contrast, dominate by *compositional abundance* — they are simply the most numerous complete-genome chromosomes recovered from this borehole — and their per-chromosome cluster-membership rates (2.3 and 1.8 clusters per genome respectively) are at or below the rest-of-universe mean.

The Omnitrophota result has direct biological reading. The majority of described Omnitrophota genomes encode host-attachment gene clusters (Seymour et al. 2023), and a hyperactive cross-replicon mobilome is consistent with the predator-/symbiont-style lifestyle inferred for the phylum: chromosomes embedded in close-contact ecological associations have repeated opportunity to acquire and exchange cargo with co-resident plasmids, integrative elements, and phages (Castelle et al. 2018, Tian et al. 2020, Wu et al. 2024). The compositional dominance of Patescibacteriota and Nanobdellota in the participant cohort is, by contrast, a measurement-level effect — these small-genome lineages are abundant and complete-genome-recoverable in this borehole, but their individual chromosomes do not carry an elevated cross-replicon cargo load. Cell-cell contact mechanisms documented in DPANN and Patescibacteriota (Johnson et al. 2024, Hamm et al. 2019) remain the plausible delivery route when transfer does occur, alongside conventional conjugation and transduction (mechanistically reviewed for archaea in Wagner et al. 2017).

### Two large lineage-restricted divergent integrative mobile elements

The 233-kbp u20424375 and the 123-kbp u29249220 are the most consequential single contigs in this study. Together they account for 289 of 791 cross-replicon clusters (37% of the cohort); both assemble as closed circles in long-read sequencing, both carry overwhelmingly uncharacterised protein content (most predicted proteins do not hit Pfam profiles at all), and both lack canonical Pfam plasmid / phage signatures. We characterise them as large divergent integrative mobile elements of a class not well-represented in current reference data.

Their host range is lineage-restricted, not broad. For u20424375, whole-genome alignment recovers exactly one recent integration host (u17976357, JAHKAP01 genus); 57% of its cross-replicon clusters involve only same-genus partners, 40% reach other Omnitrophota genera, and only 4% (7 of 195) reach beyond Omnitrophota. For u29249220, the inferred primary host (u2110955, UBA6249 genus) is in 100% of the element’s clusters; 69% (65/94) involve only UBA6249 partners, 26% reach other Omnitrophota genera, and only 5% (5 of 94) cross-phylum. Lineage-restriction is the biologically expected outcome for elements at this size: large mobile elements (>50 kb) require host-specific replication, partition, and integration machinery — Rep / iterons for copy-number control (Maddamsetti et al. 2025 establish the inverse power-law between plasmid size and copy number across 12,006 plasmids), partition systems coordinating with host nucleoid architecture, and integrase chemistry tuned to specific att sites — and these requirements constrain large-MGE host ranges to closely related lineages (Smillie et al. 2010, Suzuki et al. 2010). The cross-phylum cluster fraction we observe (4–5% per hub) is consistent with cargo-protein-mediated cluster formation across distant chromosomes (universal cargo proteins clustering across phyla without active inter-phylum transfer of the element itself) rather than active broad-host-range mobility.

The mobility apparatus on u20424375 is a Mu-class DDE transposase (Pfam Transposase_mut) sitting at the att region of the circular form; no tyrosine recombinase or serine resolvase is detectable. The element is therefore replicative-transposition-class, not a canonical conjugative ICE. Its cargo is dominated by an essentially complete bacterial big-operon r-protein cluster (31 r-protein KOs) with no rRNA genes and only four tRNAs — a profile that, to our knowledge, has no published precedent: the most translation-rich mobile genetic element previously described carries up to two r-proteins per element (Mizuno et al. 2019, across >10,000 viral genomes), and the ICEberg 3.0 catalogue of >2,000 curated ICEs (Wang et al. 2024) catalogues none with r-protein-operon cargo.

In contrast to u20424375’s Mu-class DDE chemistry, u29249220’s mobility apparatus is a tyrosine integrase (HTH_23 DNA-binding domain + integrase catalytic domain) within the 8.7-kbp integrative module at positions 103,765–112,499 — site-specific recombination of the lambda Int / ICE Int class, not replicative transposition. The two hubs therefore converge on the same population-level pattern (top two by cross-replicon involvement, both lineage-restricted within Omnitrophota, both carrying overwhelmingly uncharacterised protein content) via two distinct integration mechanisms. This argues against a single “divergent Omnitrophota mobile-element class” explanation: the cohort contains at least two distinct integrative-element families that converge on the same population-level pattern of within-genus lineage-restricted mobilisation.

The cargo is u17976357’s chromosomal r-protein region in episomal state, not extra essential cargo: the same 31 r-protein roles are present at the equivalent contiguous chromosomal locus on the two same-genus congeners (u4770637, u26960546), where they are not associated with an integrated mobile element. We therefore interpret u20424375 as a Mu-class element that mobilises a chromosomal region of u17976357 containing the big-operon ribosomal cluster, a hyper-variable polysaccharide cassette (see below), and additional accessory cargo — not as a delivery vehicle for essential rescue genes. Coverage analysis is consistent with the one-physical-copy-two-states model expected of an active integrative mobile element: chromosomal-integration + episomal coverage at backbone regions sums to ∼1.2× single-copy chromosome baseline, the small overshoot consistent with replicative transposition transiently producing extra episomal copies during the transposition cycle.

We release both contigs and their predicted proteomes as part of the dataset.

### A hyper-variable polysaccharide cassette on u20424375 mirrors the Pelagibacter HVR pattern

Within the 99.3%-identity backbone alignment of u20424375 to its u17976357 integration site, one ∼26-kb stretch (chromosomal positions 547–572 kb) is too divergent to align under the strict settings and is recovered only under more permissive realignment, at 70–86% nucleotide identity. The diverged region encodes the same proteins in the same order as the host’s chromosomal counterpart — an O-antigen / capsule / polysaccharide biosynthesis cassette including tyrosine-protein kinase Etk/Wzc, O-antigen ligase, lipopolysaccharide exporter, alginate O-acetyltransferase complex AlgI/AlgJ, multiple glycosyltransferases, and lipopolysaccharide heptosyltransferase II. Read coverage analysis on both the chromosomal and episomal cassette regions shows the same signature: the cassette is at 40–55% of single-copy backbone coverage on both the integration host and on a non-integrated congener (u26960546), indicating that the cassette is hyper-variable across the JAHKAP01 strain swarm and that ∼40–60% of element-bearing molecules in the sample carry cassette variants that do not map to either of our assembled cassettes. This is the same pattern documented for the *Pelagibacter* (SAR11) hypervariable region (Nielsen and Lui 2026a), in which a single surface-polysaccharide locus per genome undergoes rapid recombinational diversification across closely related strains. The Pelagibacter analogy is sharper still in light of Holmfeldt et al.’s (2021) “boom and burst” framing of Äspö viral dynamics: surface-polysaccharide variation is a canonical phage-escape mechanism (Labrie et al. 2010; Rodriguez-Valera et al. 2009), and a strain swarm carrying multiple cassette variants is a canonical hedge against phage burst cycles.

### *rbcL* as a chromosome-only transfer in deep groundwater

Nielsen and Lui (2026b) report lateral transfer of the carbon-fixation gene *rbcL* (RuBisCO large subunit; KEGG K01601) between Nanobdellota archaea and Patescibacteriota, in dialogue with an earlier *rbcL* LGT analysis (Jaffe et al. 2019). In KR0015B alone, an HMM scan with the K01601 model recovered 37 *rbcL* cluster representatives in the full 740,492-rep cohort, anchored on 37 complete chromosomes. Notably, none of the 37 *rbcL*-bearing clusters falls in the 791 cross-replicon cohort because no *rbcL* hit lands on a contig classified as a mobile partner.

The *rbcL*-bearing chromosomes span Patescibacteriota and Nanobdellota — the two phyla highlighted by Nielsen and Lui (2026b) — together with Altiarchaeota, Thermoproteota, Aenigmatarchaeota, Halobacteriota, Iainarchaeota, and Micrarchaeota. A protein-tree of the 37 rbcL representatives interleaves archaeal and bacterial leaves (archaeal MRCA contains 21 non-archaeal leaves; bacterial MRCA contains 18 non-bacterial leaves), but no cross-domain leaf-pairs sit as immediate sister-relatives: Form-level groupings (in the Form I/II/III/IV nomenclature of Tabita et al. 2007) remain coherent within each domain even though Forms themselves are scattered across the host tree. The pattern is consistent with multiple deep transfer events on the timescale of prokaryote-domain divergence (Battistuzzi et al. 2004) that subsequently radiated within each receiving domain, rather than ongoing same-Form transfer.

This case study reinforces the dual-catalog framing of the previous section: the cross-replicon cohort is conservative and biology-rich but not complete, and a specific well-characterised LGT story falls outside it for a structural reason. The *rbcL* transfer in KR0015B is recoverable as cross-chromosome LGT but invisible to the mobile-partner-required cross-replicon filter; it appears in the wider 957-cluster catalog (one of the 37 rbcL clusters reaches the ≥2 chromosomes-from-≥2 phyla threshold there). Nielsen and Lui (2026b) present the per-protein phylogeny in full.

### Limitations

This study covers one borehole, one assembly, one timepoint. We make no claims about temporal dynamics, between-borehole variation, or the relative rates of cross-replicon transfer compared to other environments. The cross-replicon filter requires both partners to be present and detectable in the sample at high stringency; any transfer whose mobile vehicle has been lost or has diverged below that threshold is invisible. The mobile-partner classification scheme relies on Pfam-A and geNomad reference data, both of which are biased toward well-studied phage and plasmid families; the two large divergent integrative mobile elements identified above are explicit examples of where this bias matters.

### Synthesis

In a single 9,382-contig assembly from a Fennoscandian deep-groundwater borehole, 791 cross-replicon clusters link 199 chromosomes to 367 circular mobile elements, dominated by small-genome lineages on the chromosome side (Patescibacteriota, Omnitrophota, and Nanobdellota) and by Caudoviricetes phages and conjugative plasmids on the mobile side. Two large divergent integrative mobile elements (u20424375 and u29249220) — both lineage-restricted within Omnitrophota and carrying overwhelmingly uncharacterised protein content — account for over a third of all cross-replicon clusters and lack canonical plasmid / phage signatures; the 233-kbp u20424375 carries a Mu-class DDE transposase and an essentially complete bacterial big-operon r-protein cluster as cargo, with no known precedent in the published mobile-element literature. Seven cross-replicon clusters span both domains; per-cluster phylogenies confirm that 6 of the 10 callable smoking-gun trees recover gene-tree topologies that violate the species-tree expectation. Restricting to the 791 cohort substantially under-estimates the chromosome-only LGT footprint: a parallel cross-chromosome catalog without the mobile-partner requirement contains 957 clusters, 95% of which are invisible to the cross-replicon filter (and where the *rbcL* transfer story documented by Nielsen and Lui (2026b) sits, alongside other chromosome-only transfers). 97% of cross-replicon cluster representatives carry at least one Pfam, KEGG-KO, or UniRef90 annotation, with the recognisable cargo classes — restriction-modification systems, toxin-antitoxin pairs, IS-family transposases, type IV pilus and secretion machinery — collectively accounting for the recognisable accessory-genome content. The dataset is released as a community resource for downstream characterisation of the cargo and the mobile-element classes that carry it.

## Methods

### Sample and sequencing

Sample KR0015B was collected from the Äspö Hard Rock Laboratory at 69 m below sea level. DNA extraction and Oxford Nanopore PromethION sequencing followed the protocol described in Nielsen and Lui (2026b).

### Assembly and circular-contig recovery

The PromethION reads were assembled with myloasm v0.3.0 with `-b 250` (Bloom filter size; non-default). Per-unitig circularity, length, and read coverage were read from the assembler’s `KR0015B_info.tsv` output. Coverage is taken as `Depth1` (reads aligning at ≥99% identity); the stricter `Depth2` (≥99.75%) and `Depth3` (100%) thresholds collapse to within-strain abundance and are overly conservative for an ONT environmental metagenome. Contigs flagged as `Circular = yes` and of length ≥5,000 bp were retained, yielding 9,382 circular contigs for downstream analysis.

### Gene prediction

Pyrodigal v3.6.3 (Larralde 2022; underlying Prodigal algorithm: Hyatt et al. 2010) in metagenomic mode (`-p meta -n -j 32` ; `-n` sets `force_nonsd=True`, skipping Shine-Dalgarno motif training) called genes on the 9,382 circular contigs. Output: 1,082,188 predicted proteins.

### Protein clustering

MMseqs2 (Steinegger and Söding 2017; linclust backbone: Steinegger and Söding 2018) easy-cluster was run on the predicted proteins with --min-seq-id 0.70 -c 0.80 --cov-mode 0 --cluster-mode 0 -s 7.5 --threads 32, producing 740,492 clusters (137,634 multi-member; 125,323 multi-contig).

### Cross-replicon filter

CheckM2 v1.1.0 (Chklovski et al. 2023) was run on the 1,600 assembly contigs of length ≥500 kbp (regardless of circularity); 500 kbp was our practical chromosome-candidate input threshold for this dataset, not a CheckM2 lower bound. The chromosome universe was defined as the intersection of CheckM2 high-quality (completeness ≥90%, contamination ≤5%) and circular in the long-read assembly: 237 complete genomes. We held the threshold strict. Mobile partners are circular contigs that are not chromosomes — extrachromosomal replicons (plasmids, phages, integrative elements freed in the assembly graph, and other small-circular replicating units). Operationally, the mobile-partner set comprises circular contigs of length <400 kbp — a heuristic cut below which closed circular contigs in environmental long-read assemblies are overwhelmingly plasmids or phages rather than chromosomes — plus four manually-promoted candidate jumbo phages (see below). The full size distribution across all 9,382 circular contigs is given in Supplementary Table S1.

Four marker-free ≥1 Mbp circular contigs (u30481432, u31431355, u5053247, u5294102) — each failing GTDB-Tk classification, lacking cellular markers (TIGRFAM bacterial dnaA / archaeal Cdc6/Orc1), and showing assembly multiplicity > 1 — were manually promoted into the mobile-partner set as candidate jumbo phages.

The questionable set comprises 63 circular contigs ≥400 kbp that are neither complete genomes nor jumbo-phage candidates (CheckM2 below high-quality, or no CheckM2 result) — biologically heterogeneous (candidate megaplasmids, partial chromosomes, chimeric assemblies). Each was classified by running the Pfam-A v38.1 phage panel (818 profiles) and plasmid panel (247 profiles) on its predicted proteome and assigning a per-contig call (`phage_candidate` / `plasmid_candidate` / `mixed` / `weak`); the full annotated list is Supplementary Table S2. None of the 63 contigs is used in any cross-replicon number reported here.

Each cluster was characterised by the classes of its member contigs. The cross-replicon LGT candidate cohort was defined as clusters spanning at least one chromosomal contig and at least one mobile contig; this yielded 794 candidate clusters before housekeeping filtering (see *Housekeeping filter*, below) and 791 after, involving 199 distinct chromosomes and 367 distinct mobile contigs. The chromosomal-contig set is enforced by quality and topology, not by size, which excludes incomplete chromosomal-class circular contigs from the chromosome universe.

### Chromosome taxonomy

The 199 participating chromosomes were classified with GTDB-Tk v2.7.1 (Chaumeil et al. 2020, 2022) using the GTDB R232 reference (Parks et al. 2022). One chromosome (u4168876) tripped HMMER’s 100,000-AA target-sequence cap on a 147-kAA single-frame ORF during the TIGRFAM identify step. For taxonomic placement only, proteins were re-called externally with Pyrodigal v3.6.3 (non-meta mode, `force_nonsd=True`), predictions ≥100,000 AA were dropped, and the resulting FAA was supplied to GTDB-Tk via `--genes` against the same R232 reference. All 199 chromosomes were classified, as 168 Bacteria (bac120 marker set) and 31 Archaea (ar53 marker set).

### Mobile-partner classification

geNomad v1.11.2 (Camargo et al. 2024) was run end-to-end against the v1.9 reference database with `--cleanup` on the 572 mobile-contig members of an upstream cross-class LGT filter (clusters spanning ≥2 distinct contig-size classes among chromosome / large-mobile / typical-mobile / small-mobile). The per-contig classifications were subsequently subset to the 367 mobile contigs that share at least one protein cluster with the chromosome universe. Two complementary HMM panels were built from Pfam-A v38.1 (Mistry et al. 2021) by keyword search over `Pfam-A.hmm.dat` :

- **Plasmid panel**: profiles whose name or description matched replication initiator, Rep_, RepA, RepB, RepC, Replicase, MOB_, MobA, MobC, MobL, Relaxase, VirB[0-9], T_virB, TraD, TraG, TraI, Conjug, Initiator_C, ParA, ParB → 247 profiles.
- **Phage panel**: profiles matching Phage_*, Capsid, major capsid, Terminase, Portal, Tail_*, tail tape, tape measure, Baseplate, head morphogenesis, integrase_*, prophage, Viral_*, holin, endolysin, polyhedrin → 818 profiles.

A small chromosomal-marker panel (TIGR00362, bacterial dnaA; TIGR02928, archaeal Cdc6/ Orc1 — both from the GTDB-Tk-bundled TIGRFAM HMMs) was run as a misassembly screen.

The three panels were searched against the geNomad-output protein FASTA on the 367 mobile contigs using `hmmsearch` from HMMER v3.4 (Eddy 2011) with `-E 1e-5 --cpu 16` . Per-contig integrated classification rules:

- **plasmid (strong)** if (≥2 distinct Pfam plasmid HMMs hit) OR (geNomad plasmid score ≥0.7 AND ≥1 Pfam plasmid HMM hit)
- **virus (strong)** if (≥2 distinct Pfam phage HMMs hit) OR (geNomad virus score ≥0.7 AND ≥1 Pfam phage HMM hit)
- **both** if both strong criteria are met simultaneously
- **plasmid (single line of evidence)** if geNomad plasmid score ≥0.7 but no Pfam plasmid HMM hits, OR if a single Pfam plasmid HMM hits but geNomad plasmid score is <0.7; analogous for virus (single line of evidence) with the phage panel
- **unclassified** if no signal of either class

The 77-vs-37 disagreement on the 133 geNomad-unresolved contigs (Results) tracks the lower plasmid recall reported for geNomad’s neural-network classifier (plasmid MCC 0.778 vs virus MCC 0.953; Camargo et al. 2024), which reflects phage-skewed training data. The Pfam-A keyword-derived plasmid panel above recovers the missed cohort at modest computational cost.

### Phylogenetic confirmation of LGT

Per-cluster maximum-likelihood phylogenies were built for the 12-cluster smoking-gun cohort (the 7 cross-domain clusters and the 5 housekeeping-filter-passing cross-phylum clusters with the broadest phylum span — ≥3 distinct bacterial phyla on participating chromosomes). Member protein sequences were extracted from the cross-replicon protein FASTA, deduplicated by sequence id (Pyrodigal output occasionally duplicates the bare-id header line), aligned with mafft v7.525 (Katoh and Standley 2013) in L-INS-i mode (`--localpair --maxiterate 1000`), trimmed with BMGE v2.0 (Criscuolo and Gribaldo 2010) using `-t AA -m BLOSUM30` (default entropy and gap thresholds), and supplied to IQ-TREE 3 (Wong et al. 2026) with `-T 16 --mset LG,WAG,JTT --mrate I,G -B 1000` for ultrafast bootstrap branch support (Hoang et al. 2018). The model panel was restricted to the three matrices and two rate models that dominate ModelFinder selection on small bacterial/ archaeal protein alignments of the kind we are scoring; broadening the panel was not expected to alter topology at the level of the disruption-score read-out. Phylum-disruption scores were computed by, for each phylum *P* present in a tree, taking the smallest clade containing all *P*-leaves and counting non-*P* leaves with assigned phylum within that clade; the per-tree score is the sum of these counts divided by the tree’s total leaves with assigned phylum. The same logic was used to evaluate domain-clean splits in cross-domain trees (archaeal vs bacterial leaves form reciprocal clades vs intermixed). Treefiles and per-tree disruption tables are released as Supplementary Table S4.

### Cross-chromosome companion catalog

A complementary catalog was built by removing the mobile-partner requirement from the cross-replicon filter and retaining clusters whose members reach ≥2 distinct complete-genome chromosomes from ≥2 distinct phyla. The catalog was then filtered against the housekeeping-marker panel described below. The output (957 clusters; Supplementary Table S3) is reported alongside the 791 cohort to bound the chromosome-only LGT landscape that the cross-replicon filter intentionally excludes.

### Housekeeping-marker filter

Universal single-copy housekeeping proteins (ribosomal proteins, RNA polymerase subunits, translation factors, key replication and repair enzymes) are deeply conserved across phyla and can retain ≥70% AA identity in pairwise comparisons across the deepest splits of the tree, particularly among slow-evolving lineages. They therefore can clear the cross-replicon clustering threshold not because of recent transfer but because of deep orthology — and their inclusion would inflate the cohort with deep-ortholog signal masquerading as recent LGT. To exclude this class, both the cross-replicon and the cross-chromosome cohorts were filtered against the GTDB-Tk bac120 + ar53 single-copy-marker HMM panels. The marker lists (16 PFAM + 152 TIGRFAM, deduplicated across the two panels) were loaded at runtime from the canonical GTDB-Tk source (`gtdbtk.config.common`) rather than hand-typed, and the corresponding HMMs were taken from the GTDB-Tk R232 reference distribution. Cluster representatives were searched with HMMER v3.4 `hmmsearch -E 1e-10` against the combined marker panel. Any cluster whose representative scored a hit was dropped. The filter removed 3 of 794 (0.4%) clusters from the cross-replicon cohort — all three were r-protein or EF-Tu clusters carried by u20424375 — and 27 of 984 (2.7%) clusters from the cross-chromosome catalog. The full list of dropped clusters with the matched marker is released as Supplementary Table S9.

### Per-phylum cross-replicon enrichment test

Each chromosome’s cross-replicon cluster-membership count was computed by summing membership across the 791 cluster set; each phylum’s distribution of per-genome counts was tested against the rest of the 212 GTDB-classified chromosomes with a one-sided Mann-Whitney U test (`scipy.stats.mannwhitneyu(…, alternative=’greater’)`), Benjamini-Hochberg FDR-corrected across phyla. Chromosomes whose only cross-replicon participation was through housekeeping clusters (and that therefore have a post-filter count of zero) were retained in the test universe with their zero counts. The 25 chromosomes in the universe that are GTDB-unclassified were excluded from the test. The Mann-Whitney test has minimal statistical power for phyla represented by ≤2 chromosomes; results for these small phyla are released in Supplementary Table S7 as descriptive counts only, and the headline per-genome enrichment claims rest on the well-sampled phyla (Omnitrophota n = 77, Patescibacteriota n = 71, Nanobdellota n = 19). Per-phylum statistics are released as Supplementary Table S7.

### Integration mapping for u20424375

u20424375 was aligned against all 237 KR0015B chromosomes with minimap2 v2.28 (Li 2018) under `-x asm5` (intended for sequences ≤1% divergent) to identify high-identity recent integration sites; u17976357 was the only chromosome onto which ≥80% of the element aligned at ≥98% identity. To recover diverged regions within the integrated copy, the same pair was realigned under `-x asm20` (permissive for sequences up to ∼20% divergent), which exposed the ∼26-kb hyper-variable polysaccharide cassette at 70–86% identity within an otherwise 99.3%-identity backbone.

### Chimera screen on the divergent hubs

The two large divergent integrative mobile elements (u20424375, u29249220) were screened for assembly chimeras with a base-resolution read-mapping test on the KR0015B Oxford Nanopore reads. The three flowcells of KR0015B PromethION reads were mapped with minimap2 v2.28 (`-x map-ont -a -t 32` ; Li 2018) against a 7-contig reference comprising the two hubs (each doubled head-to-tail to allow reads spanning the circular origin to map across the seam) and five high-leverage chromosomal partners; alignments were sorted with samtools v1.20 (Danecek et al. 2021). For each contig we computed per-position read depth, end-count (reads whose alignment starts or ends at the position), and soft-clip pile-up (reads with a soft-clip operation ≥100 bp at the position), summed across all three BAMs; for the doubled hubs the second-half mapping was folded onto canonical positions before signal calculation. A genuine assembly chimera produces a depth cliff (depth → 0 at a single position because reads cannot physically span an artefactual concatenation), accompanied by a localised end-pile-up and soft-clip spike; a real multi-copy biological feature (rRNA operon, IS-element cluster, integrative cargo block) instead produces a depth dip across a few kilobases without depth approaching zero. Both hubs show no depth-cliff signature: the minimum smoothed depth in any flagged window on u20424375 and u29249220 is 36× and 25× respectively, well above the cliff threshold and consistent with real circular elements. The screen additionally identified the 8.7-kbp embedded integrative module on u29249220 reported in Results. Per-position TSVs and per-contig verdicts are released as Supplementary Table S8.

### *rbcL* census

The K01601 KEGG-ortholog HMM (KofamScan v1.3.0 profile distribution; Aramaki et al. 2020) was scanned with `hmmsearch -E 1e-5` (HMMER v3.4) over both the 791 cross-replicon cluster representatives and the full 740,492-rep cohort, after filtering predicted proteins of length ≥100 kAA that exceed HMMER’s per-target pipeline limit. Hits were intersected with the cluster.tsv table to determine cluster-level membership; contigs were classified as HQ chromosome / mobile partner / large non-HQ circular via the same contig classes used in the cross-replicon filter. Across the 37 clusters, member contigs split into 37 complete chromosomes and 36 large (≥400 kbp) non-complete circulars; no cluster contains a mobile-partner contig. Per-rbcL-cluster contig partitioning is in Supplementary Table S5.

### Functional annotation of cross-replicon cluster representatives

The 791 cluster representatives were annotated against three independent sources:

1. **Pfam-A v38.1** (Mistry et al. 2021): `hmmsearch -E 1e-5 --cpu 64` .
2. **KofamScan** with the KEGG ortholog profile panel (Aramaki et al. 2020): `exec_annotation -E 1e-5 --cpu 64` .
3. **UniRef90**: mmseqs easy-search --min-seq-id 0.3 -e 1e-5 -c 0.5 --cov-mode 0 .

Best hit per representative per source was retained. The aggregated table (`per_cluster_annotation.tsv`) keys on cluster representative and is released as Supplementary Table S6. InterProScan was attempted but the bundled HMM datasets in this build (Pfam-A 381 kB, TIGRFAM 510 kB) are stub data and yielded only 4 / 791 hits; the full IPS data archive (∼80 GB) was not downloaded for this run because the direct Pfam-A search above already covered Pfam-side annotation at full resolution.

## Code and data availability

The KR0015B Oxford Nanopore reads and assembled contigs are available under BioProject PRJNA1457649. Pipeline scripts and supplementary tables S1–S9 will be deposited on Zenodo upon acceptance, with an accompanying README documenting the per-script inputs/outputs and the contents of each supplementary table.

