## Supplementary material for "Omnitrophota dominate per-genome cross-replicon lateral gene transfer in a Fennoscandian deep-groundwater metagenome": README.md

# Supplementary tables — LGT paper

Each table is a tab-separated values file (`.tsv`) keyed by the column described below.

| File | Description |
|---|---|
| `S1_circular_contig_size_distribution.tsv` | Per-unitig length, depth, multiplicity for all 9,382 circular contigs in the KR0015B assembly (myloasm `info.tsv` filtered to `Circular = yes`, `Length >= 5 kbp`). |
| `S2_questionable_circular_contigs.tsv` | 63 large (>=400 kbp) circular contigs that are neither high-quality chromosomes nor jumbo-phage candidates, with per-contig Pfam-based phage/plasmid classifications. None are used in any cross-replicon number in the paper. |
| `S3_cross_chromosome_hgt.tsv` | The wider cross-chromosome catalog (957 clusters; cross-domain + cross-phylum, mobile-partner requirement dropped). |
| `S4_smoking_gun_topology.tsv` | Per-tree phylum-disruption scores and per-leaf metadata for the 12-cluster smoking-gun cohort. |
| `S5_rbcl_landscape.tsv` | Per-cluster contig partitioning for the 37 *rbcL* (K01601) clusters in the full 740,492-rep cohort. |
| `S6_per_cluster_annotation.tsv` | Best Pfam-A / KofamScan-KO / UniRef90 hit per cluster representative across the 791 cross-replicon cohort. |
| `S6_annotation_coverage.txt` | Per-source annotation-coverage summary (counts behind Figure 8A). |
| `S7_phylum_enrichment_test.tsv` | Per-phylum Mann-Whitney U one-sided test of per-genome cross-replicon participation against the rest of the chromosome universe (BH-FDR-corrected). |
| `S8_chimera_screen.tsv` | Per-position read-mapping signal (depth, end-count, soft-clip pile-up) and chimera-verdict for u20424375, u29249220, and five high-leverage chromosomal partners. |
| `S8_chimera_screen_summary.txt` | Human-readable verdict summary for S8. |
| `S9_housekeeping_filter_dropped_clusters.tsv` | The 27 clusters dropped by the GTDB-Tk bac120 / ar53 housekeeping-marker filter, with the strongest matching marker per cluster and a flag indicating whether the cluster also belonged to the 794-pre-filter cross-replicon cohort (the 3 cross-replicon drops are the ones flagged). |

The full pipeline (scripts, intermediate files, marker lists, and the cluster-rep FASTA) will be deposited on Zenodo at acceptance.
